# Graph theory analysis reveals an assortative pain network vulnerable to attacks

**DOI:** 10.1101/2023.03.08.531580

**Authors:** Chong Chen, Adrien Tassou, Valentina Morales, Grégory Scherrer

**Author notes:** **Corresponding authors:** Dr. Chong Chen, Dr. Grégory Scherrer.

## Abstract

The neural substrate of pain experience has been described as a dense network of connected brain regions. However, the connectivity pattern of these brain regions remains elusive, precluding a deeper understanding of how pain emerges from the structural connectivity. Here, we use graph theory to systematically characterize the architecture of a comprehensive pain network, including both cortical and subcortical brain areas. This structural brain network consists of 49 nodes denoting pain-related brain areas, linked by edges representing their relative incoming and outgoing axonal projection strengths. Sixty-three percent of brain areas in this structural pain network share reciprocal connections, reflecting a dense network. The clustering coefficient, a measurement of the probability that adjacent nodes are connected, indicates that brain areas in the pain network tend to cluster together. Community detection, the process of discovering cohesive groups in complex networks, successfully reveals two known subnetworks that specifically mediate the sensory and affective components of pain, respectively. Assortativity analysis, which evaluates the tendency of nodes to connect with other nodes with similar features, indicates that the pain network is assortative. Finally, robustness, the resistance of a complex network to failures and perturbations, indicates that the pain network displays a high degree of error tolerance (local failure rarely affects the global information carried by the network) but is vulnerable to attacks (selective removal of hub nodes critically changes network connectivity). Taken together, graph theory analysis unveils an assortative structural pain network in the brain processing nociceptive information, and the vulnerability of this network to attack opens up the possibility of alleviating pain by targeting the most connected brain areas in the network.

## Introduction

Pain is a complex, multidimensional, and subjective experience. Neuroimaging and neurophysiological studies have shown that noxious stimuli activate an extensive network of cortical brain areas, which has been termed the *pain matrix* ^1–4^. Historically, this functional pain network has been divided into sensory-discriminative and cognitive-affective systems ^5–7^. The sensory-discriminative system, which includes the lateral thalamus and primary and secondary somatosensory cortices (SI and SII, respectively), is thought to process information related to nociceptive inputs, including intensity, localization, and quality ^8,9^. The cognitive-affective system, which comprises brain regions such as the anterior insula (AI) and anterior cingulate cortex (ACC), is believed to mediate psychological aspects of pain ^4,10–12^. The concept of the *pain matrix* implies that there is no single “pain center” in the brain, and that the activity pattern of this functional network could serve as a reliable and objective indicator of painful experience, including in pathophysiological pain states in which pain may occur in the absence of any nociceptive stimulus ^13–15^. However, recent studies suggest that the *pain matrix* activation is a response to salient sensory stimuli rather than perceptual qualities unique to pain ^16–18^. Therefore, it’s crucial to include additional brain areas, especially subcortical brain regions within ascending and descending pain pathways that participate in nociceptive information transmission, processing and modulation, to study pain at the network level.

Graph theory, a branch of mathematics concerning the formal description and analysis of graphs, provides a powerful method to characterize network structure and function ^19^. Using graph-theoretical tools, pain networks can be modeled as a set of functional or structural interactions. In these models, nodes, denoting brain areas, are linked by edges, representing structural or functional connections between them. In previous studies, researchers in the pain field have built functional pain networks using data from noninvasive neuroimaging and neurophysiology techniques, detecting altered resting network topology in patients with chronic pain disorders ^20–23^. However, functional pain networks lack reciprocal connection information between different brain regions ^24,25^, and are typically unable to include deep brain structures due to technical limitations. Thus, the structural topological properties of pain networks, and how these properties support multidimensional pain remain elusive. To overcome these limitations, we sought to construct a structural pain network that comprises both superficial and deep brain areas involved in pain perception, and uses anatomical reciprocal axon connections as edges preserving the direction information.

We utilized the Allen Mouse Brain Connectivity Atlas, a comprehensive open-access online database containing high-resolution images of traced axonal projections from defined mouse brain regions and cell types ^26,27^, to generate a directed and weighted pain network. In this network, we included all brain areas that, according to our literature search, may play a role in pain. Graph theory analysis of this structural pain network revealed that: 1) 63% of brain areas in the pain network share reciprocal connections; 2) clustering analysis indicated that brain areas in the pain network tend to cluster together; 3) community detection successfully identified and separated the two known systems for the sensory and affective components of pain, respectively; 4) the assortativity coefficient reflects an assortative pain network; and 5) robustness evaluation showed that the pain network displays a high degree of error tolerance but is vulnerable to attacks.

## Results

### Brain areas included in the structural pain network

Precisely which regions constitute the *pain matrix* has yet to be conclusively and consistently defined ^4^. Thus, to create a comprehensive brain network for pain, we conducted a literature search in an effort to include all brain areas that have been reported to participate in pain perception (Table S1). In addition, considering that the pontine nuclei (PG) and inferior olivary nuclei (IO) mediate communication between the cerebral cortex and cerebellum, which critically contribute to pain processing ^28^, we included both the PN and IO in this network. In total, this pain network comprises 49 major brain subdivisions based on the Allen Reference Ontology ^29^. Among them, 18 brain structures are located in the cerebrum (CH), 21 in the brain stem (BS), and 10 in the cerebellum (CB).

### Noxious stimulation activates all brain areas in the network

To confirm that brain areas in this structural pain network participate in pain perception, we used mutant TRAP2 (*Fos*^CreERT2^);Ai14 mice to genetically label with tdTomato neurons that are active during nociception, using noxious pinprick as a stimulus (Figure 1A). Neurons expressing tdTomato were widely distributed throughout the brain (Figure 1B). Although the number of TRAPed neurons in each brain area varied, we detected tdTomato-positive neurons in every brain area in this structural pain network (Figure 1C). Among the CH group, the retrosplenial cortex (RSP), motor cortex (MO), and anterior cingulate cortex (ACC) displayed extensive tdTomato labeling (Figure 1C and D), consistent with their critical roles in pain perception. From the BS group, the paraventricular nucleus of the thalamus (PVT), nucleus raphe pontis (RPO) and parabrachial nucleus (PB), all of which are intensively investigated in the pain field, showed the highest density of tdTomato-positive neurons (Figure 1C and D). Besides, we also observed a wide distribution of tdTomato-positive neurons in the cerebellar cortex and nuclei (Figure. 1C and D), in line with accumulating evidence suggesting that the cerebellum participates in pain processing ^30^.

**Figure 1.**
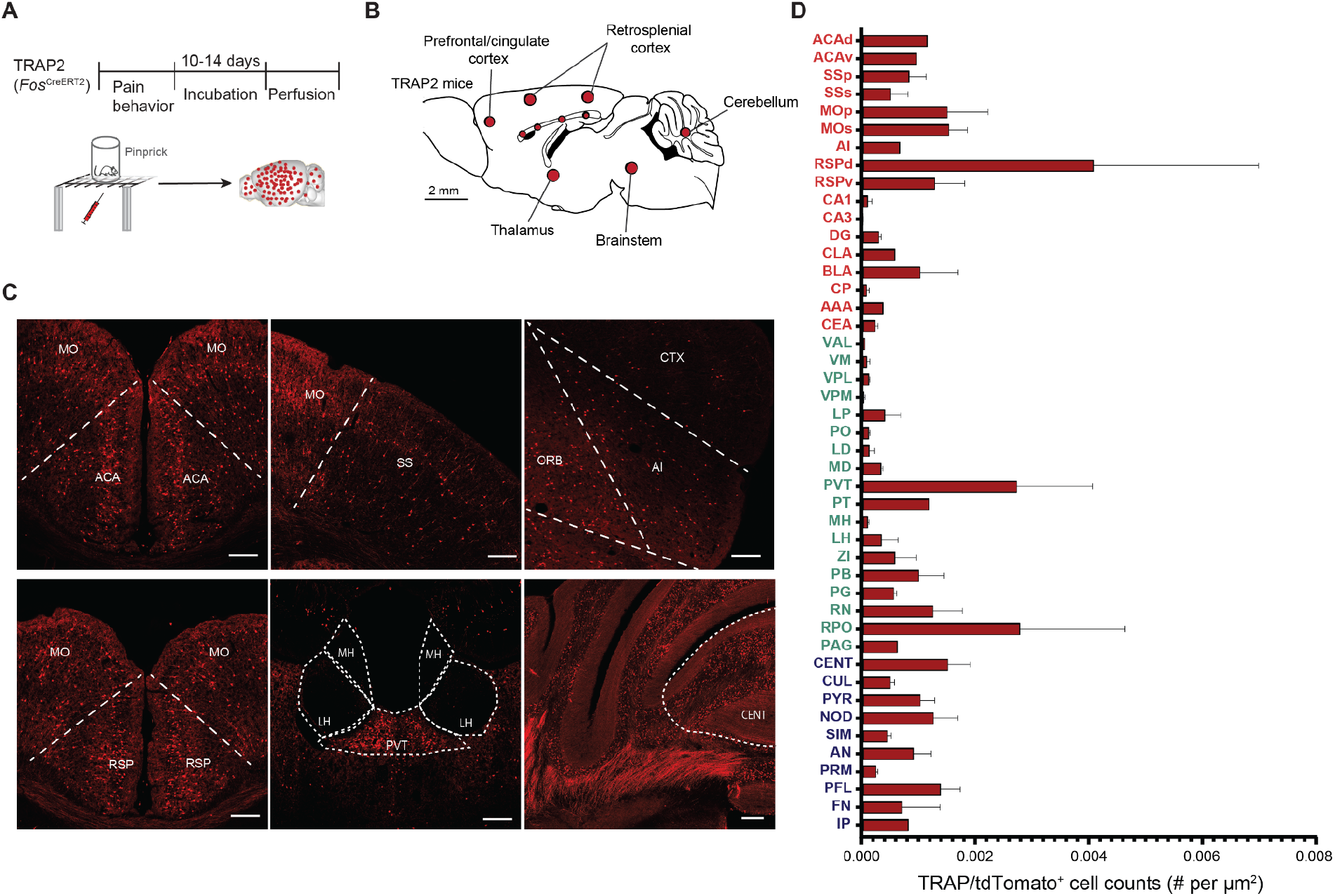
Noxious pinprick stimulation activates neurons in diverse brain areas. (A) Experimental timeline to label neurons active during noxious pinprick stimulation. (B) Overview of tdTomato-expressing neurons in various brain areas. The size of each circle (red) was scaled based on the density of labeled neurons. (C) Representative photomicrographs of tdTomato-expressing neurons in several brain areas. Scale bars, 200 μm. (D) Quantification of tdTomato-expressing neurons across brain areas in the pain network (n = 3).

### Ipsilaterally dominated and distance-dependent connectivity

To build a structural pain network, we queried the normalized projection volumes, defined as the total volume of segmented pixels (EGFP signal) in the target normalized by the injection site volume, of each brain area to other regions of the pain network from the Allen Mouse Brain Connectivity Atlas using the API (http://help.brain-map.org/display/mouseconnectivity/API). We restricted our analysis to wildtype mice and only included data from mutant mice when data from wildtype mice was not available for 6 brain structures (for details, please see the Materials and Methods). Connectivity strengths between brain areas in this pain network span a greater than 10^5^-fold range (Figure 2), suggesting that the quantitative physical connections between brain areas in the pain network must be considered in order to understand, interpret and discriminate its activity patterns during pain perception ^26,31^. In addition, the connectivity matrix shows prevalent bilateral projections between brain areas in the network, with generally stronger ipsilateral projections than contralateral (total normalized projection volumes are 3.5:1 between the ipsilateral and contralateral hemispheres). Of all possible connections above the minimal true positive level of 10^-4 26^, 59% project ipsilaterally, while 41% project contralaterally.

**Figure 2.**
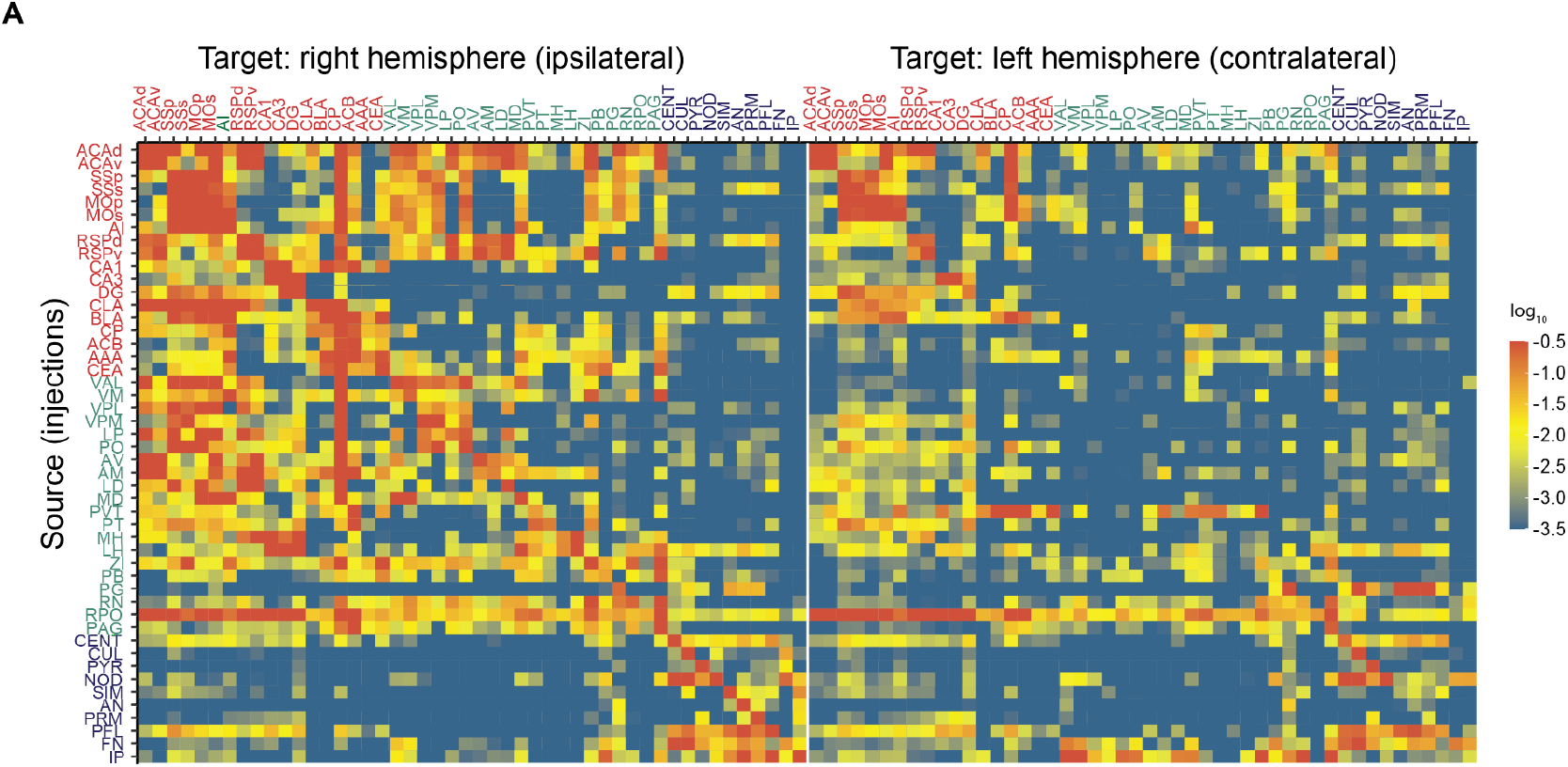
Connectivity matrix of brain areas in the pain network. (A) Each row shows the quantitative projection signals from one of the 49 brain areas to other brain areas (in columns) in the right (ipsilateral) and left (contralateral) hemispheres. Brain areas are displayed in ontological order. Color maps indicate log_10_-transformed projection strength. All values less than 10^-3.5^ are shown in blue to minimize false positives. All values greater than 10^-0.5^ are shown in red to reduce the dominance of projection signals in certain large regions ^26^.

Brain areas in the CH and BS groups are extensively inter- and intraconnected, whereas CB brain areas show sparse intraconnection (Figure 3A). The limited connection between cerebellar subareas is consistent with the lobular anatomical organization of the cerebellum ^32,33^. In addition, the lack of direct connections between the CB and brain areas in the CH and BS groups reflects the indirect nature of cortico-cerebellar and cerebello-cortical connectivity ^34,35^. Furthermore, connection strengths between brain areas in the pain network display a hemisphere-correlated distance dependence (Figure 3B and C). In the ipsilateral hemisphere, connection strength between two brain areas decreases as the distance between them increases. Whereas in the contralateral hemisphere, we found no clear correlation (Figure 3B and C). These findings mirror the anatomy of the somatosensory pathways, in which each side of the body is represented contralaterally in the brain.

**Figure 3.**
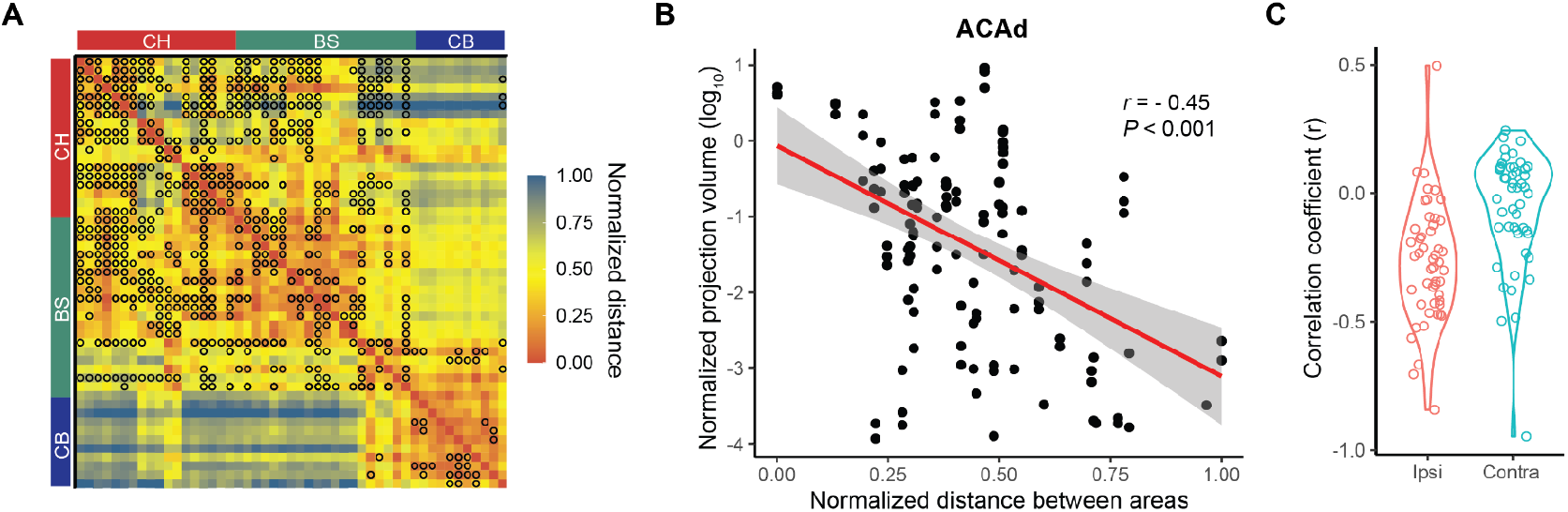
Distance-dependent connections between brain areas in the pain network. (A) Distance matrix of all brain areas (ipsilateral) in the pain network. Color map indicates the normalized distance between brain areas based on injection coordinates. Black circle indicates a projection strength larger than 0.01. Brain areas are displayed in ontological order and divided into anatomical groups: CH, BS or CB. (B) Scatter plot of the projection strength from the dorsal part of the anterior cingulate area, dorsal part (ACAd) to other brain areas in the pain network as a function of the distance between them. Data points were fit by linear regression (red line). (C) Violin plot of the correlation coefficient between projection strength and distance of each brain area to other brain areas (ipsilateral and contralateral) in the pain network.

### Homogenous connection pattern between brain areas in the network

Nodes with similar connection patterns in a network tend to exhibit similar functionality ^36^. Thus, we analyzed the similarity in connection patterns of different brain areas in the pain network. We compared the similarity of outgoing projections originating from two brain areas (source correlation) and the similarity of incoming projections terminating in these two areas (target correlations). Heatmaps of both source (Figure 4, left) and target correlations (Figure 4, right) indicated two clusters: one belonging to the CH and BS groups, and the other belonging to the CB group (Figure 3A). Among these two clusters, a large number of brain areas in the pain network showed both strong incoming and outgoing projection correlations (Figure 3A), indicating a homogenous connection pattern between brain areas in the network.

**Figure 4.**
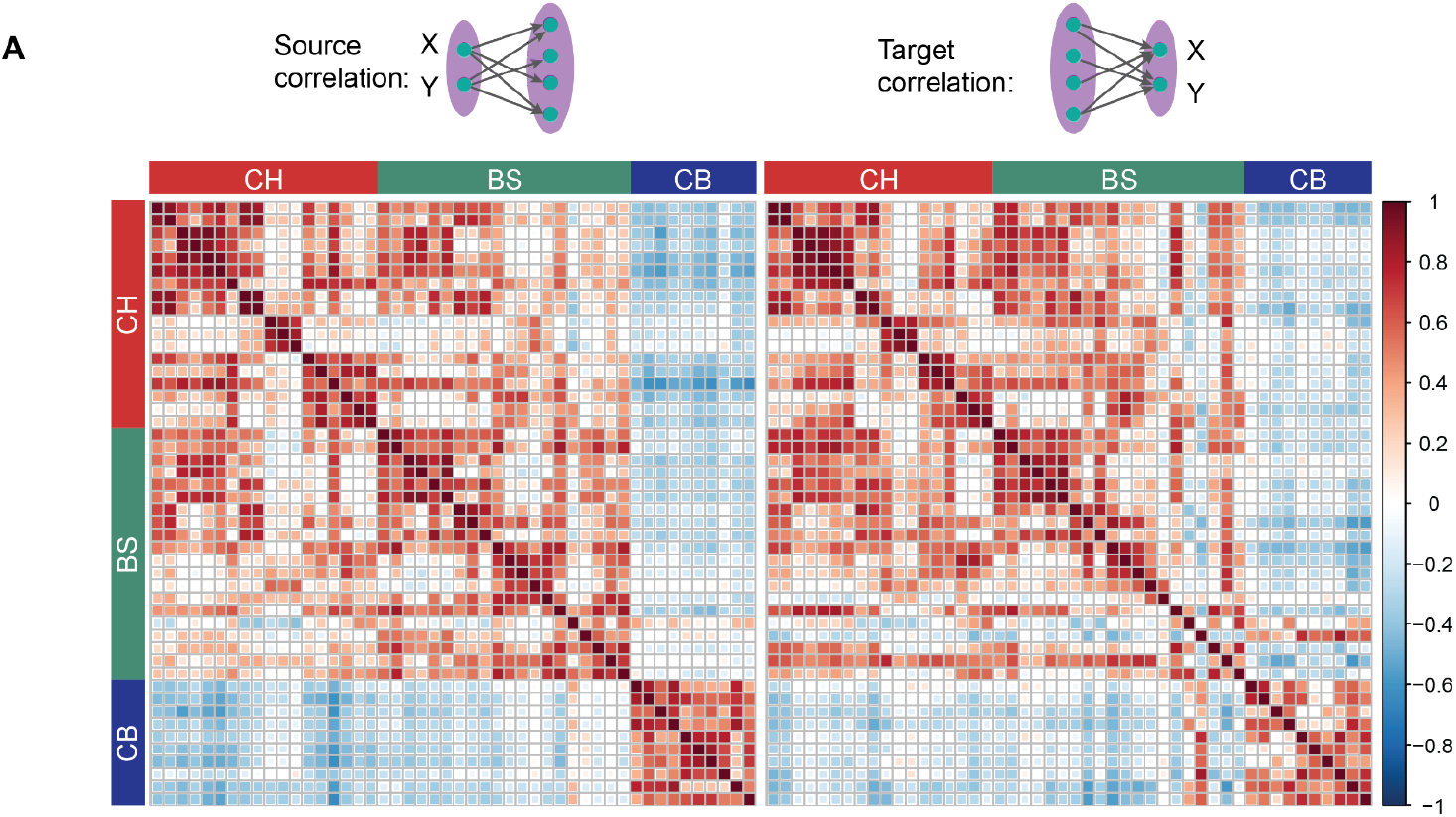
Correlation coefficients of projection strengths between areas. (A) Comparison of the correlation coefficients of projection strengths between brain areas in the pain network, defined as the common source for projection to other regions (left) and as the common target of projection from other regions (right). Color maps indicate the correlation coefficient value. Brain areas are displayed in ontological order and divided into three anatomical groups: CH, BS or CB.

### Overview of the pain network architecture

To investigate the topological properties of this structural pain network, we generated a directed and weighted network according to the axonal connections between brain areas (Figure 5A). The pain network contains 49 nodes and 632 edges, yielding a network density (indicative of how efficiently a network transmits information) of 27%. The pain network has a diameter (the shortest distance between the two most distant nodes) of 7 nodes. Network diameter determines how quickly information could spread through a network ^37^. In addition, 63% of nodes in the pain network share reciprocal connections, compared to 26% in a generated random network with the same density (Table 2). These results indicate that brain areas in the pain network are densely interconnected.

**Figure 5.**
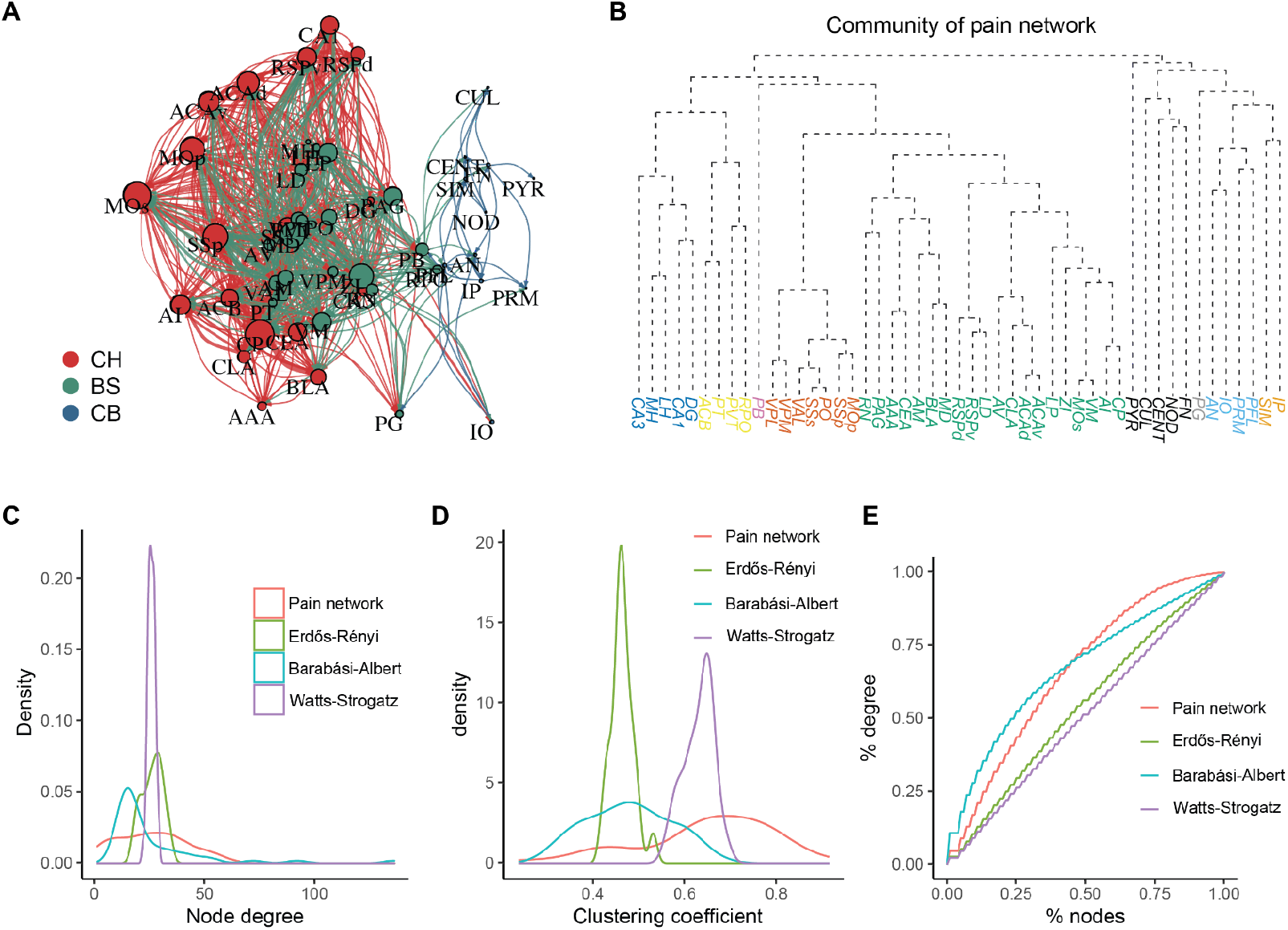
Topological properties of the pain network. (A) Graphical layout of the pain network. The position of each vertex corresponds to its injection site coordinates scaled to a two-dimensional plane. Node size is scaled according to the degree centrality of each node. Color indicates the group to which a vertex belongs to. (B) Community detection of the pain network using random walks. (C) Vertex degree distribution of the pain network, compared against the corresponding Erdős-Rényi random graph model, the Watts-Strogatz small-world model and the Barabási-Albert model. (D) Similar to (B), comparison of the vertex clustering coefficient of each network model. (E) Lorenz concentration curve showing the degree distribution among the four network models.

### Hub nodes in the pain network

Hub nodes, which connect with many other nodes of a network, typically contribute more to network function than relatively isolated nodes ^19^. To identify the hub nodes of this pain network, we conducted centrality analysis using three different methods (Table 1 and Table S2). Degree centrality, which assesses the number of edges belonging to each node, revealed that the caudoputamen (CP), secondary motor cortex (MOs) and mediodorsal nucleus of thalamus (MD) share the highest number of direct connections with other nodes of the pain network. Closeness centrality measures the average shortest distance from each node to each other node, while betweenness centrality determines the likelihood that a node sits between other nodes in networks. Notably, the parabrachial nuclei (PB) display both the highest closeness centrality and betweenness centrality in the pain network, supporting the notion that PB is a hub for pain ^38^.

**Table 1.** Top three vertices in different centrality analysis.

| Centrality | Top1 | Top2 | Top3 |
| --- | --- | --- | --- |
| Degree | CP | MOs | MD |
| Closeness | PB | MOp | CEA |
| Betweenness | PB | CEA | ZI |

### Communities for sensory and affective pain

To test whether graph theory analysis could detect the known subsystems for the sensory-discriminative and affective-motivational components of pain, we conducted community detection using the Walktrap algorithm ^39^. In total, this algorithm detected 13 communities in the pain network (Figure 5B and Table S3). Remarkably, based only on the projection data, this algorithm clearly separated the respective networks critical for the sensory and affective components of pain ^5–7^. In addition to the somatosensory cortex and lateral thalamic nuclei, which play a crucial role in sensory pain processing ^40–43^, the Walktrap algorithm included the primary motor cortex (M1), but not the secondary motor cortex (M2), in this community. For the community mediating the affective dimension of pain, other than the well-established ACC, agranular insular area (AI), mediodorsal nucleus of the thalamus (MD) and BLA ^11,44–47^, the algorithm further identified the periaqueductal gray (PAG), retrosplenial area (RSP), zona incerta (ZI) and red nucleus (RN) as belonging to this community.

### Pain network appears assortative

To further understand the global structure of this pain network, we compared its topological properties with three mathematical models of networks with the same density: the Erdős-Rényi random graph model ^37^, Watts-Strogatz small-world model ^48^ and Barabási-Albert model ^49^ (scale-free model; Figure 5C-D and Table 2). Graph theory analysis revealed that the pain network has the highest mean clustering coefficient (0.65) among these four networks (Table 2). The node degree distribution of a small-world network model resembles that of the pain network (Figure 5B); however, its clustering coefficient distribution aligns poorly (Figure 5C). The degree distribution of the pain network is right-skewed compared to the random graph model and small-world model, indicating that a substantial portion of the edges are concentrated on a small number of highly connected nodes (Figure 5B). The scale-free model also shows a right-skewed degree distribution, but does not closely fit the pain network.

**Table 2.** Topological properties comparison between networks.

| Modle | Reciprocity | Clustering | Assortativity |
| --- | --- | --- | --- |
| Pain network | 0.63 | 0.65 | 0.14 |
| Erdős–Rényi | 0.26 | 0.46 | -0.03 |
| Watts-Strogatz | 1 | 0.48 | 0.55 |
| Barabási–Albert | 0 | 0.63 | -0.07 |

To test whether the pain network is assortative or disassortative, we conducted assortativity analysis. Biological networks tend to be disassortative ^50,51^. In these networks, a strong effective repulsion between highly connected nodes increases the specificity of functional modules and stability against random network error ^52,53^. However, the pain network displays a positive assortativity coefficient (Table 2), indicating that, unlike most biological networks, the pain network is assortative.

### Error and attack tolerance of the pain network

Many complex systems, including the World-Wide Web, the Internet, social networks and the cells, display a high-degree of error tolerance at the expense of attack vulnerability ^54^. To assess how the pain network would respond to random failure and attack, we removed a fraction (*f*) of either nodes randomly or hub nodes from the pain network ^54^. The removal of any node generally increases the distance between the remaining nodes, as it can eliminate some paths that contribute to the interconnectedness of the system ^54^. By examining the mean distance (the average length of the shortest path between any two nodes) of remaining nodes, we found that the pain network displays a high degree of error tolerance: the ability of their nodes to communicate is barely affected (Figure 6A), even at a high failure rate (removing 20% nodes in the pain network randomly). However, removing hub nodes from the pain network substantially increases the mean distance, indicating that the pain network is vulnerable to attacks (Figure 6A).

**Figure 6.**
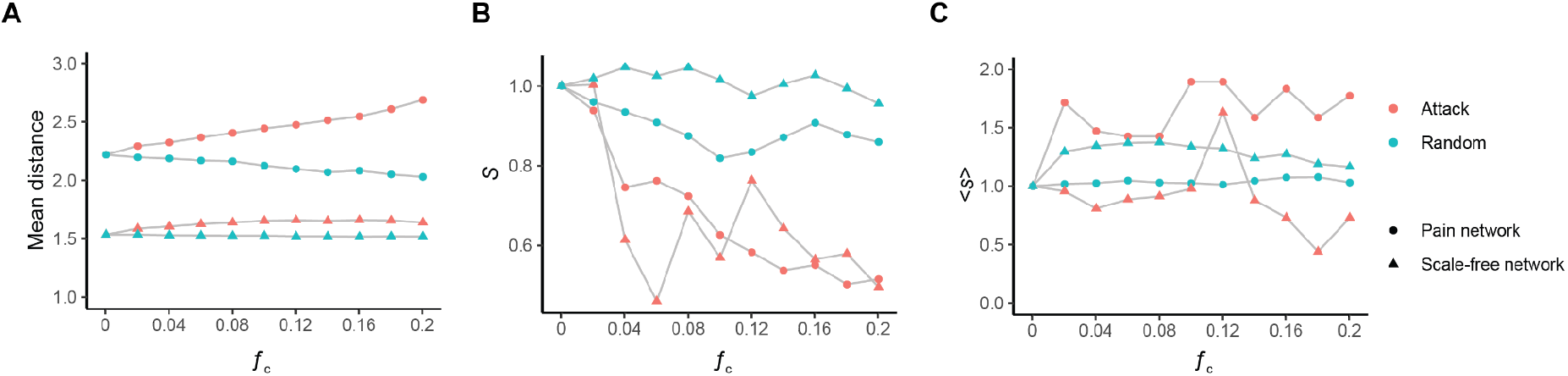
Pain network displays tolerance against both random failure and attack. (A) Change in the mean distance of the pain network as a function of the fraction *f* of the removed random nodes (Random, cyan) or hub nodes (Attack, red). (B) The relative size of the largest cluster *S* as a function of the fraction *f* of the removed random nodes (Random, cyan) or hub nodes (Attack, red) for the pain network (circle) and a scale-free network with the same density (triangle). (C) The average size of the isolated clusters as a function of the fraction *f* of the removed random nodes (Random, cyan) or hub nodes (Attack, red) for the pain network (circle) and a scale-free network with the same density (triangle).

Removing nodes from a network can induce network fragmentation, because clusters of nodes whose links to the system may be cut off ^54^. To assess the impact of random failures and attacks on network fragmentation of the pain network, we measured the size of the largest cluster, *S*, shown as a fraction of the total system size, when a fraction, *f* of the nodes is removed either randomly or in an attack. We found that, when removing nodes randomly, as *f* increases, *S* remains stable. In contrast, *S* decreases dramatically during attacks, behaving similarly to a disassortative scale-free network with the same number of nodes and edges (Figure 6B). Nevertheless, the pain network behaves differently than a scale-free network when assessing the average size of isolated clusters (all clusters except the largest one), <s>. We found that <s> remains stable for the pain network during random failure while it increases during attacks (Figure 6C), suggesting that the remaining nodes form new clusters. Taken together, these results indicate that the pain network, like other complex systems, displays a high degree of tolerance against random failure, and the pain network is vulnerable to attacks, but responds differently than typical scale-free networks.

## Discussion

Using projection data from the Allen Brain Connectivity Atlas, we built a structural pain network and analyzed it using graph theory. We found that brain areas in this pain network display extensive reciprocal connections, and these connections show ipsilateral distance dependence. Based solely on projection data, community detection clearly identified and separated the two systems for the sensory versus affective component of pain (Figure 5B and Table S3). Compared with three common random models, the pain network has a positive assortativity coefficient (Table 2), indicating that this pain network is assortative. Finally, robustness analysis indicated that the pain network shows a high degree of error tolerance but is vulnerable to attacks (Figure 6). The attack vulnerability of the pain network opens up the possibility of alleviating pain by targeting hub brain nodes.

### Structure-function relations of the pain network

Physical connection strength between brain areas in this structural pain network varies widely (Figure 2). This is in line with the diverse functional connectivity strength between brain areas determined by pairwise correlation analysis of their activity during pain ^23,55^. However, a high functional correlation between brain areas does not necessarily entail a strong physical connection ^56,57^, because additional factors contribute to functional correlation, such as the quantity, strength (e.g., synaptic efficiency), and dynamic modulation (e.g., short- and long-term synaptic plasticity) of the physical connections ^58^. Network-level factors also play a role. For example, two brain areas without any direct connection can show highly correlated activity via one or more intermediate brain areas ^59,60^. How physical connections lead to the functional correlation between brain areas during pain, and the mechanisms by which structural and functional connectivities change during chronic pain are interesting subjects for future study.

Several brain areas in the cerebral cortex, including the ACC, motor cortex and somatosensory cortex show strong interhemispheric connections (Figure 2). Previous studies have shown that interhemispheric communication of the dorsolateral prefrontal cortex (DLPFC) influences pain tolerance and discomfort by modulating interhemispheric inhibition ^61,62^. In addition, unilateral optogenetic stimulation of pyramidal neurons in the somatosensory cortex can prevent or reduce mechanical hypersensitivity bilaterally ^63^. Thus, additional studies are still needed to elucidate the role of cortical interhemispheric connections in pain processing.

### Hubs of the pain network

Hub nodes facilitate information integration by occupying a highly connected and functionally central position in the network ^64^. Hubs can be identified using several forms of centrality analysis, such as degree-based, strength-based and path-based ^65^. However, all of these measures have limitations when identifying the hubs of functional networks generated using data from brain imaging techniques ^65,66^. We conducted topological centrality analysis of our structural pain network using degree, betweenness and closeness centrality (Table 1 and Table S2). Degree centrality analysis indicated that the CP, MOs and MD are the most highly connected nodes in this pain network, consistent with their critical roles in pain perception ^67–70^. Betweenness and closeness centrality (path-based measures) identified the PB, which has been recognized as a sensory hub for both pain and aversion ^38,71^, as the topological center of the pain network.

### Communities for multi-dimensional pain

Pain is a multidimensional experience with sensory-discriminative and affective-motivational components ^1,6,7,72^. The sensory-discriminative component determines the spatiotemporal characteristics and qualia of pain, while the affective-motivational component mediates pain unpleasantness ^73^. Although there remains debates as to the separability of these two components ^72^, two pain systems, the lateral and medial, have been proposed to contribute to the sensory and affective dimensions of pain, respectively. Remarkably, our community detection of the pain network clearly separated these two systems using only the physical projection data (Figure 5B and Table S3). Importantly, our analysis suggested several brain areas in the network that may contribute differently to the sensory or affective component of pain. For example, the red nucleus (RN) and zona incerta (ZI) were detected in the medial pain system, pointing to a role for these brain areas in affective pain (Figure 5B and Table S3). Interestingly, our community detection suggests that the primary motor cortex (MOp) contributes to the sensory component of pain, while the MOs processes the affective component (Figure 5B and Table S3). Motor cortex stimulation (MCS), primarily of the MOp, has been used clinically to treat neuropathic pain ^74,75^. Thus, it may prove interesting to investigate the effects of MOs simulation.

Further, community detection revealed several previously unidentified communities (Figure 5B and Table S3). One intriguing community comprises the hippocampal formation (CA1, CA3 and DG) and the habenular nuclei (MH and LH). The hippocampus critically contributes to learning and memory ^76,77^, while the lateral habenula plays a role in aversive learning and memory ^78,79^. Pain, particularly chronic pain, shares several commonalities with learning and memory, and has been characterized as a “painful memory” ^80–82^. Thus, this novel community could prove an exciting target for future investigations aiming to elucidate the role of learning and memory in pain, especially in pain catastrophizing.

### Pain network assortativity and robustness

Network topology determines both the efficiency ^83–85^ and strength of the entire system ^54,86^. In this study, we compared the topology of this structural pain network to three random network models with identical density: the Erdős-Rényi random graph model, Watts-Strogatz small-world model and Barabási-Albert scale-free model. None of these models perfectly fits all characteristics of the pain network, which shows a right-skewed degree distribution (Figure 5C) and the highest clustering coefficient (Figure 5D).

The pain network shows a positive assortativity coefficient (Table 2), indicating that it is assortative. Assortative networks are rare among complex biological networks, which tend to be disassortative ^50,51^. In contrast, assortative networks abound in social science, in which individuals tend to bond with others who share similar characteristics ^87,88^. Assortativity facilitates cooperation in social activities ^89^ and promotes information distribution ^90^. Thus, the positive assortativity coefficient of the pain network indicates that brain areas in the network share similarity in connectivity, which may facilitate pain processing.

In addition, our analysis indicates that this assortative pain network shows a high degree of error tolerance to random failures (Figure 6). The error tolerance of the pain network provides a possible structural network mechanism underlying the intractability observed in some cases of chronic pain ^91^. However, unlike assortative networks which are resilient, the pain network is vulnerable to attacks, behaving similarly to disassortative scale-free networks ^50,54^.

### Attack vulnerability of the pain network

Most social and technological networks display an unexpected degree of error tolerance. For instance, we rarely experience global network outages despite frequent rooter issues. However, the error tolerance of these networks comes at the expense of attack survivability: removing the most connected node substantially changes the connectivity of a network. Attacking search engines nowadays, such as Google, would significantly affect our ability to surf and locate information on the web.

Interestingly, the assortative pain network is vulnerable to attacks (Figure 6). The attack vulnerability of the pain network indicates that the pain network could be damaged by targeting hub brain nodes. Indeed, deep brain stimulation in the striatum ^92^, motor cortex ^93,94^ and thalamus ^95,96^, the densely connected nodes in the pain network (Table 1 and Table S2), has been used for pain control in clinics. While such treatments confer particularly effective pain relief ^97^, they have primarily been empirical. Our study provides a network-level explanation for the effectiveness of these applications, potentially paving the way for the development of new therapies that target hub nodes in the pain network to enhance their efficacy further.

In conclusion, our graph theory analysis revealed an assortative pain network in the brain for pain processing. The pain network displays a high degree of tolerance against random failure but is vulnerable to attacks. This structural pain network reflects the hierarchical organization of the functional pain network, and may underlie the diverse and persistent activity patterns during acute and chronic pain. Combined with functional neuroimaging, neurophysiology techniques and machine learning, graph theory analysis of the pain network could provide a novel way to objectively diagnose, at the system level, both acute and chronic pain. Most importantly, this deeper understanding of the pain network could lead to novel therapeutic methods for pain management.

## Acknowledgements

This project was supported by grants from the National Institutes of Health R01NS106301 (G.S.), R21DA049241 (G.S.), R01NS118504 (G.S.), the McKnight Endowment Fund for Neuroscience (G.S), the Brain Research Foundation (G.S.), the New York Stem Cell Foundation (G.S.). G.S. is a New York Stem Cell Foundation – Robertson Investigator. We thank Dr. Nicole Mercer Lindsay for reading the manuscript and J. Blair for manuscript editing.

## Author contributions

C.C. conceptualized and designed the project. C.C. performed the analysis. A.T. and V.M. collected and analyzed the histology data from TRAP2 mice. C.C. wrote the manuscript with input from all authors. G.S. acquired funding.

## Data and code availability

The R code used to query projection data from Allen Brain Institute, pain network analysis and visualizations is provided at https://github.com/chenchong446337/pain_netwrok. The projection data from each brain area to others in the pain network are also included.

## Materials and Methods

### Animals

All procedures followed animal care guidelines approved by the Institutional Animal Care and Use Committee of the University of North Carolina at Chapel Hill, and by the International Association for the Study of Pain. Mice were housed at a maximum of 5 per cage and maintained on a 12-hr light/dark cycle in a temperature-controlled environment with ad libitum access to food and water. TRAP2 (*Fos*^CreERT2^) mice were purchased from Jackson Laboratory (Stock #: 030323).

### Determining pain-related brain areas

To generate a comprehensive pain network, we conducted a literature search via Google Scholar (https://scholar.google.com/) and Pubmed (https://pubmed.ncbi.nlm.nih.gov/), aiming to include all brain areas reportedly involved in pain. We defined brain areas according to the Allen Reference Atlas ontology. Search keywords included “pain,” “nociception,” “chronic pain” and “neuropathic pain.” If a general brain region was reported to participate in pain, its functional subdivisions were included as individual nodes of the pain network.

### Drugs

4-OHT (H6278, Sigma) was prepared in absolute ethanol and Kolliphor EL (C5135, Sigma) and administered intraperitoneally (50 mg/kg).

### Pinprick to TRAP neurons active during pain

Mice were habituated in a cylinder for at least 30 min before the experiment. We then delivered noxious pinprick (25-G needle) stimuli to the plantar surface of the hindpaw of TRAP2 (*Fos*^CreERT2^);Ai14 mice. In total, we administered 20 stimuli at 30 s intervals. After stimulation, mice were allowed to remain in the cylinders for an additional 2 hr before receiving 4-OHT (50 mg/kg) injection subcutaneously. After 4-OHT injection, mice were allowed to remain in the cylinders for another 1 hr before being returned to their homecages. Two weeks later, we perfused the mice and dissected the brains to verify tdTomato expression.

### Histology

Animals were transcardially perfused with phosphate-buffered saline (PBS) followed by 4% formaldehyde in PBS. Brains were then dissected, post-fixed in 4% formaldehyde, and cryoprotected in 30% sucrose. Tissues were then frozen in Optimum Cutting Temperature compound (OCT; product code: 4583, Tissue Tek) and sectioned using a cryostat (Leica). Brains were sectioned at 40 μm and stored in PBS at 4°C if used immediately. For longer-term storage, tissue sections were placed in glycerol-based cryoprotectant solution and stored at −20°C. For in situ hybridization, tissues were sectioned at 14 μm, collected on Superfrost Plus slides (catalog: 22-037-246, Fisher Scientific) and stored at −80°C.

### Imaging and cell counting

For imaging, sections were mounted on slides using Fluoromount-G (SouthernBiotech) and left at room temperature overnight for proper polymerization. Images were collected using a Carl Zeiss LSM 780 confocal microscope and processed with ImageJ (NIH). Cells expressing tdTomato were counted both manually and automatically using the “analyze particle” toolbox.

### Projection data query from Allen Brain Atlas

After determining which brain areas participate in the pain network, we queried the projection data of each brain area via the Allen Brain Atlas API (Application Programming Interface) using R 4.0.2 software (The R Project for Statistical Computing). Projection data of most brain areas in the pain network were queried from wildtype mice. For six brain areas (CLA, VPL, MH, PG, RPO, IO) with no experiments performed in wildtype mice, we used the projection data from mutant mice. If multiple experiments were returned for one brain area, the projection intensity to other brain areas was calculated by averaging the projection intensity from all experiments. The injection coordinates of each brain area were collected and used to calculate the distance between brain areas (Figure 2).

### Visualization

All figures were generated using R 4.0.2. The position of each vertex was based on the injection coordinates for each brain area. We first calculated the distance between each brain area, then scaled it to two-dimensional coordinates using the “cmdscale” function from the “stats” package. The edge width of each vertex was determined by the projection strength. For better visualization, we performed a log transformation and scaled it down 5 times.

### General features analysis

Using the projection data of each brain area, we created an adjacency list and converted it into a directed graph using the “graph.data.frame” function from the “igraph” package ^98^ (https://igraph.org) running on R. To exclude weak and/or spurious connections, brain areas with projection values of < 0.01 were not considered to be connected ^26^. To visualize the network in 2D, the distances between brain areas are embedded into a 2D plane using multidimensional scaling.

Using the “igraph” package, we calculated the graph-theoretical metrics to evaluate the following general network properties: density, diameter and reciprocity. The “edge_density” function was used to calculate the density of the pain network. The density of a graph is the ratio of the number of edges to the number of possible edges, 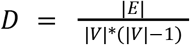. We used the “diameter” function to calculate the diameter of the pain network. The diameter of a graph is the length of the longest geodesic. We used the “reciprocity” function to measure the proportion of nodes in the pain network that share reciprocal connections.

### Network centrality analysis

Using the functions from the “igraph” package, we evaluated the centrality of the pain network with three common measures: degree centrality, closeness centrality and betweenness centrality (Table 1 and Table S2). We used the “degree” function to determine the number of adjacent edges for each node in the network. The pain network we constructed is a directed network; thus, we analyzed the total degree, which is the sum of the in- and out-degree of each node. We used the “closeness” function to measure closeness centrality, an indicator of how close each node is to every other node in a network. Finally, we used the “betweenness” function to measure node betweenness, defined by 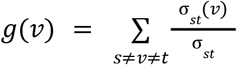, where σ_*st*_ is the total number of shortest paths from node *s* to node *t* and σ_*st*_(*v*) is the number of those paths that pass through *v.* Betweenness indicates the likelihood that a node sits between each pair of other nodes in the network.

### Community detection

We used the “cluster_walktrap” function to detect the communities of the pain network (Figure 5B and Table S3). This function uses the Walktrap community finding algorithm ^39^ to identify densely connected subgraphs.

### Random network models

We generated three random network models to compare against the pain network. We used the “erdos.renyi.game” function to generate a random network with an identical number of nodes and edge density. Next, we used the “sample_smallworld” function to create a small-world network by setting the argument “size” equal to the number of node in the pain network, “nei” equal to the average edges of each node has of the pain network, and the *p*(wiring probability) at 0.05. Finally, we generate a scale-free network using the “sample_pa” function. The argument “n” equals the number of nodes in the pain network, “power” at 0.5, using the “psumtree-multiple” algorithm.

### Robustness analysis

We assessed the error tolerance of the pain network against random failure and attack by continuously evaluating its diameter while removing a small fraction *f* of the nodes ^54^. To test tolerance against random failure, we removed nodes at random; to measure tolerance against attack, we specifically removed hub nodes. For each value of *f*, we repeated this process 100 times.

To evaluate the fragmentation process while removing nodes from the pain network, we measured the size of the largest cluster, *S*, shown as a fraction of the total system size, when a fraction *f* of the nodes were removed either at random or in an attack ^54^. We also measured the changes in average size <*s*> for all clusters except the largest, when a fraction *f* of the nodes were removed either at random or in an attack mode. For each *f*, we repeated this process 100 times.

## Reference

1. Melzack, R. (1999). From the gate to the neuromatrix. Pain Suppl 6, S121–S126.

2. Iannetti, G.D., and Mouraux, A. (2010). From the neuromatrix to the pain matrix (and back). Exp. Brain Res. 205, 1–12.

3. Ingvar, M. (1999). Pain and functional imaging. Philos. Trans. R. Soc. Lond. B Biol. Sci. 354, 1347–1358.

4. Tracey, I., and Mantyh, P.W. (2007). The cerebral signature for pain perception and its modulation. Neuron 55, 377–391.

5. Albe-Fessard, D., Berkley, K.J., Kruger, L., Ralston, H.J., 3rd, and Willis, W.D., Jr (1985). Diencephalic mechanisms of pain sensation. Brain Res. 356, 217–296.

6. Clark, W.C. (1969). Sensory-decision theory analysis of the placebo effect on the criterion for pain and thermal sensitivity. J. Abnorm. Psychol. 74, 363–371.

7. Fernandez, E., and Turk, D.C. (1992). Sensory and affective components of pain: separation and synthesis. Psychol. Bull. 112, 205–217.

8. Andersson, J.L., Lilja, A., Hartvig, P., Långström, B., Gordh, T., Handwerker, H., and Torebjörk, E. (1997). Somatotopic organization along the central sulcus, for pain localization in humans, as revealed by positron emission tomography. Exp. Brain Res. 117, 192–199.

9. Kenshalo, D.R., Jr, and Isensee, O. (1983). Responses of primate SI cortical neurons to noxious stimuli. J. Neurophysiol. 50, 1479–1496.

10. Royce, G.J., and Mourey, R.J. (1985). Efferent connections of the centromedian and parafascicular thalamic nuclei: an autoradiographic investigation in the cat. J. Comp. Neurol. 235, 277–300.

11. Meda, K.S., Patel, T., Braz, J.M., Malik, R., Turner, M.L., Seifikar, H., Basbaum, A.I., and Sohal, V.S. (2019). Microcircuit Mechanisms through which Mediodorsal Thalamic Input to Anterior Cingulate Cortex Exacerbates Pain-Related Aversion. Neuron 102, 944–959.e3.

12. Vogt, B.A. (2005). Pain and emotion interactions in subregions of the cingulate gyrus. Nat. Rev. Neurosci. 6, 533–544.

13. Borsook, D., Sava, S., and Becerra, L. (2010). The pain imaging revolution: advancing pain into the 21st century. Neuroscientist 16, 171–185.

14. Cecchi, G.A., Huang, L., Hashmi, J.A., Baliki, M., Centeno, M.V., Rish, I., and Apkarian, A.V. (2012). Predictive dynamics of human pain perception. PLoS Comput. Biol. 8, e1002719.

15. Wager, T.D., Atlas, L.Y., Lindquist, M.A., Roy, M., Woo, C.-W., and Kross, E. (2013). An fMRI-based neurologic signature of physical pain. N. Engl. J. Med. 368, 1388–1397.

16. Salomons, T.V., Iannetti, G.D., Liang, M., and Wood, J.N. (2016). The “Pain Matrix” in Pain-Free Individuals. JAMA Neurol. 73, 755–756.

17. Legrain, V., Iannetti, G.D., Plaghki, L., and Mouraux, A. (2011). The pain matrix reloaded: a salience detection system for the body. Prog. Neurobiol. 93, 111–124.

18. Iannetti, G.D., Salomons, T.V., Moayedi, M., Mouraux, A., and Davis, K.D. (2013). Beyond *metaphor: contrasting mechanisms of social and physical pain*. Trends Cogn. Sci. 17, 371–378.

19. Bullmore, E., and Sporns, O. (2009). Complex brain networks: graph theoretical analysis of structural and functional systems. Nat. Rev. Neurosci. 10, 186–198.

20. Balenzuela, P., Chernomoretz, A., Fraiman, D., Cifre, I., Sitges, C., Montoya, P., and Chialvo, D.R. (2010). Modular organization of brain resting state networks in chronic back pain patients. Front. Neuroinform. 4, 116.

21. Liu, J., Zhao, L., Li, G., Xiong, S., Nan, J., Li, J., Yuan, K., von Deneen, K.M., Liang, F., Qin, W., et al. (2012). Hierarchical alteration of brain structural and functional networks in female migraine sufferers. PLoS One 7, e51250.

22. Mansour, A., Baria, A.T., Tetreault, P., Vachon-Presseau, E., Chang, P.-C., Huang, L., Apkarian, A.V., and Baliki, M.N. (2016). Global disruption of degree rank order: a hallmark of chronic pain. Sci. Rep. 6, 34853.

23. Kaplan, C.M., Schrepf, A., Vatansever, D., Larkin, T.E., Mawla, I., Ichesco, E., Kochlefl, L., Harte, S.E., Clauw, D.J., Mashour, G.A., et al. (2019). Functional and neurochemical disruptions of brain hub topology in chronic pain. Pain 160, 973–983.

24. Wiberg, M. (1992). Reciprocal connections between the periaqueductal gray matter and other somatosensory regions of the cat midbrain: a possible mechanism of pain inhibition. Ups. J. Med. Sci. 97, 37–47.

25. Diao, Y., Cui, L., Chen, Y., Burbridge, T.J., Han, W., Wirth, B., Sestan, N., Crair, M.C., and Zhang, J. (2018). Reciprocal Connections Between Cortex and Thalamus Contribute to Retinal Axon Targeting to Dorsal Lateral Geniculate Nucleus. Cereb. Cortex 28, 1168–1182.

26. Oh, S.W., Harris, J.A., Ng, L., Winslow, B., Cain, N., Mihalas, S., Wang, Q., Lau, C., Kuan, L., Henry, A.M., et al. (2014). A mesoscale connectome of the mouse brain. Nature 508, 207–214.

27. Kuan, L., Li, Y., Lau, C., Feng, D., Bernard, A., Sunkin, S.M., Zeng, H., Dang, C., Hawrylycz, M., and Ng, L. (2015). Neuroinformatics of the Allen Mouse Brain Connectivity Atlas. Methods 73, 4–17.

28. D’Angelo, E. (2018). Physiology of the cerebellum. Handb. Clin. Neurol. 154, 85–108.

29. Wang, Q., Ding, S.-L., Li, Y., Royall, J., Feng, D., Lesnar, P., Graddis, N., Naeemi, M., Facer, B., Ho, A., et al. (2020). The Allen Mouse Brain Common Coordinate Framework: A 3D Reference Atlas. Cell 181, 936–953.e20.

30. Moulton, E.A., Schmahmann, J.D., Becerra, L., and Borsook, D. (2010). The cerebellum and pain: passive integrator or active participator? Brain Res. Rev. 65, 14–27.

31. Markov, N.T., Ercsey-Ravasz, M.M., Ribeiro Gomes, A.R., Lamy, C., Magrou, L., Vezoli, J., Misery, P., Falchier, A., Quilodran, R., Gariel, M.A., et al. (2014). A weighted and directed interareal connectivity matrix for macaque cerebral cortex. Cereb. Cortex 24, 17–36.

32. Stoodley, C.J., and Schmahmann, J.D. (2010). Evidence for topographic organization in the cerebellum of motor control versus cognitive and affective processing. Cortex 46, 831–844.

33. Miall, R.C. (2013). Cerebellum: Anatomy and Function. In Neuroscience in the 21st Century: From Basic to Clinical, D. W. Pfaff, ed. (Springer New York), pp. 1149–1167.

34. Brodal, P., and Bjaalie, J.G. (1997). Chapter 13 Salient anatomic features of the cortico-ponto-cerebellar pathway. In Progress in Brain Research, C. I. De Zeeuw, P. Strata, and J. Voogd, eds. (Elsevier), pp. 227–249.

35. Kelly, R.M., and Strick, P.L. (2003). Cerebellar loops with motor cortex and prefrontal cortex of a nonhuman primate. J. Neurosci. 23, 8432–8444.

36. White, J.G., Southgate, E., Thomson, J.N., and Brenner, S. (1986). The structure of the nervous system of the nematode Caenorhabditis elegans. Philos. Trans. R. Soc. Lond. B Biol. Sci. 314, 1–340.

37. Newman, M. (2010). Networks: An Introduction (Oxford University Press).

38. Chiang, M.C., Bowen, A., Schier, L.A., Tupone, D., Uddin, O., and Heinricher, M.M. (2019). Parabrachial Complex: A Hub for Pain and Aversion. J. Neurosci. 39, 8225–8230.

39. Pons, P., and Latapy, M. (2006). Computing Communities in Large Networks Using Random Walks. Journal of Graph Algorithms and Applications 10, 191–218. 10.7155/jgaa.00124.

40. De Ridder, D., Vanneste, S., Smith, M., and Adhia, D. (2022). Pain and the Triple Network Model. Front. Neurol. 13, 757241.

41. Liu, Y., Latremoliere, A., Li, X., Zhang, Z., Chen, M., Wang, X., Fang, C., Zhu, J., Alexandre, C., Gao, Z., et al. (2018). Touch and tactile neuropathic pain sensitivity are set by corticospinal projections. Nature 561, 547–550.

42. Ab Aziz, C.B., and Ahmad, A.H. (2006). The role of the thalamus in modulating pain. Malays. J. Med. Sci. 13, 11–18.

43. Terrier, L.-M., Hadjikhani, N., and Destrieux, C. (2022). The trigeminal pathways. J. Neurol. 10.1007/s00415-022-11002-4.

44. Corder, G., Ahanonu, B., Grewe, B.F., Wang, D., Schnitzer, M.J., and Scherrer, G. (2019). An amygdalar neural ensemble that encodes the unpleasantness of pain. Science 363, 276–281.

45. Yuan, W., Dan, L., Netra, R., Shaohui, M., Chenwang, J., and Ming, Z. (2013). A pharmaco-fMRI study on pain networks induced by electrical stimulation after sumatriptan injection. Exp. Brain Res. 226, 15–24.

46. Yang, J.-W., Shih, H.-C., and Shyu, B.-C. (2006). Intracortical circuits in rat anterior cingulate cortex are activated by nociceptive inputs mediated by medial thalamus. J. Neurophysiol. 96, 3409–3422.

47. Dong, W.K., Ryu, H., and Wagman, I.H. (1978). Nociceptive responses of neurons in medial thalamus and their relationship to spinothalamic pathways. J. Neurophysiol. 41, 1592–1613.

48. Watts, D.J., and Strogatz, S.H. (1998). Collective dynamics of “small-world” networks. Nature 393, 440–442.

49. Broder, A., Kumar, R., Maghoul, F., Raghavan, P., Rajagopalan, S., Stata, R., Tomkins, A., and Wiener, J. (2000). Graph structure in the Web. Computer Networks 33, 309–320.

50. Maslov, S., and Sneppen, K. (2002). Specificity and stability in topology of protein networks. Science 296, 910–913.

51. Maslov, S., and Sneppen, K. (2002). Protein interaction networks beyond artifacts. FEBS Lett. 530, 255–256.

52. Newman, M.E.J. (2003). Mixing patterns in networks. Phys. Rev. E Stat. Nonlin. Soft Matter Phys. 67, 026126.

53. Song, C., Havlin, S., and Makse, H.A. (2006). Origins of fractality in the growth of complex networks. Nat. Phys. 2, 275–281.

54. Albert, R., Jeong, H., and Barabasi, A.L. (2000). Error and attack tolerance of complex networks. Nature 406, 378–382.

55. Napadow, V., LaCount, L., Park, K., As-Sanie, S., Clauw, D.J., and Harris, R.E. (2010). Intrinsic brain connectivity in fibromyalgia is associated with chronic pain intensity. Arthritis Rheum. 62, 2545–2555.

56. Uddin, L.Q. (2013). Complex relationships between structural and functional brain connectivity. Trends Cogn. Sci. 17, 600–602.

57. Honey, C.J., Sporns, O., Cammoun, L., Gigandet, X., Thiran, J.P., Meuli, R., and Hagmann, P. (2009). Predicting human resting-state functional connectivity from structural connectivity. Proc. Natl. Acad. Sci. U. S. A. 106, 2035–2040.

58. Citri, A., and Malenka, R.C. (2008). Synaptic plasticity: multiple forms, functions, and mechanisms. Neuropsychopharmacology 33, 18–41.

59. Steinmetz, N.A., Zatka-Haas, P., Carandini, M., and Harris, K.D. (2019). Distributed coding of choice, action and engagement across the mouse brain. Nature 576, 266–273.

60. Douglas, R.J., Koch, C., Mahowald, M., Martin, K.A., and Suarez, H.H. (1995). Recurrent excitation in neocortical circuits. Science 269, 981–985.

61. Graff-Guerrero, A., González-Olvera, J., Fresán, A., Gómez-Martín, D., Méndez-Núñez, J.C., and Pellicer, F. (2005). Repetitive transcranial magnetic stimulation of dorsolateral prefrontal cortex increases tolerance to human experimental pain. Brain Res. Cogn. Brain Res. 25, 153–160.

62. Sevel, L.S., Letzen, J.E., Staud, R., and Robinson, M.E. (2016). Interhemispheric Dorsolateral Prefrontal Cortex Connectivity is Associated with Individual Differences in Pain Sensitivity in Healthy Controls. Brain Connect. 6, 357–364.

63. Xiong, W., Ping, X., Ripsch, M.S., Chavez, G.S.C., Hannon, H.E., Jiang, K., Bao, C., Jadhav, V., Chen, L., Chai, Z., et al. (2017). Enhancing excitatory activity of somatosensory cortex alleviates neuropathic pain through regulating homeostatic plasticity. Sci. Rep. 7, 12743.

64. van den Heuvel, M.P., and Sporns, O. (2013). Network hubs in the human brain. Trends Cogn. Sci. 17, 683–696.

65. Fornito, A., Zalesky, A., and Bullmore, E.T. eds. (2016). Chapter 5 - Centrality and Hubs. In Fundamentals of Brain Network Analysis (Academic Press), pp. 137–161.

66. Power, J.D., Schlaggar, B.L., Lessov-Schlaggar, C.N., and Petersen, S.E. (2013). Evidence for hubs in human functional brain networks. Neuron 79, 798–813.

67. Barceló, A.C., Filippini, B., and Pazo, J.H. (2012). The striatum and pain modulation. Cell. Mol. Neurobiol. 32, 1–12.

68. Brasil-Neto, J.P. (2016). Motor Cortex Stimulation for Pain Relief: Do Corollary Discharges Play a Role? Front. Hum. Neurosci. 10, 323.

69. Mo, J.-J., Hu, W.-H., Zhang, C., Wang, X., Liu, C., Zhao, B.-T., Zhou, J.-J., and Zhang, K. (2019). Motor cortex stimulation: a systematic literature-based analysis of effectiveness and case series experience. BMC Neurol. 19, 48.

70. Bushnell, M.C., Duncan, G.H., Hofbauer, R.K., Ha, B., Chen, J.I., and Carrier, B. (1999). Pain perception: is there a role for primary somatosensory cortex? Proc. Natl. Acad. Sci. U. S. A. 96, 7705–7709.

71. Roeder, Z., Chen, Q., Davis, S., Carlson, J.D., Tupone, D., and Heinricher, M.M. (2016). Parabrachial complex links pain transmission to descending pain modulation. Pain 157, 2697–2708.

72. Talbot, K., Madden, V.J., Jones, S.L., and Moseley, G.L. (2019). The sensory and affective components of pain: are they differentially modifiable dimensions or inseparable aspects of a unitary experience? A systematic review. Br. J. Anaesth. 123, e263–e272.

73. Auvray, M., Myin, E., and Spence, C. (2010). The sensory-discriminative and affective-motivational aspects of pain. Neurosci. Biobehav. Rev. 34, 214–223.

74. Cha, M., Um, S.W., Kwon, M., Nam, T.S., and Lee, B.H. (2017). Repetitive motor cortex stimulation reinforces the pain modulation circuits of peripheral neuropathic pain. Sci. Rep. 7, 7986.

75. Garcia-Larrea, L., and Peyron, R. (2007). Motor cortex stimulation for neuropathic pain: From phenomenology to mechanisms. Neuroimage 37 Suppl 1, S71–S79.

76. Jarrard, L.E. (1993). On the role of the hippocampus in learning and memory in the rat. Behav. Neural Biol. 60, 9–26.

77. Bird, C.M., and Burgess, N. (2008). The hippocampus and memory: insights from spatial processing. Nat. Rev. Neurosci. 9, 182–194.

78. Tomaiuolo, M., Gonzalez, C., Medina, J.H., and Piriz, J. (2014). Lateral Habenula determines long-term storage of aversive memories. Front. Behav. Neurosci. 8, 170.

79. Wang, D., Li, Y., Feng, Q., Guo, Q., Zhou, J., and Luo, M. (2017). Learning shapes the aversion and reward responses of lateral habenula neurons. Elife 6. 10.7554/eLife.23045.

80. Price, T.J., and Inyang, K.E. (2015). Commonalities between pain and memory mechanisms and their meaning for understanding chronic pain. Prog. Mol. Biol. Transl. Sci. 131, 409–434.

81. Mansour, A.R., Farmer, M.A., Baliki, M.N., and Apkarian, A.V. (2014). Chronic pain: the role of learning and brain plasticity. Restor. Neurol. Neurosci. 32, 129–139.

82. Sandkühler, J. (2000). Learning and memory in pain pathways. Pain 88, 113–118.

83. Pastor-Satorras, R., and Vespignani, A. (2001). Epidemic spreading in scale-free networks. Phys. Rev. Lett. 86, 3200–3203.

84. Leskovec, J., Adamic, L.A., and Huberman, B.A. (2007). The dynamics of viral marketing. ACM Trans. Web 1, 5 – es.

85. Wu, F., Huberman, B.A., Adamic, L.A., and Tyler, J.R. (2004). Information flow in social groups. Physica A: Statistical Mechanics and its Applications 337, 327–335.

86. Tu, Y. (2000). How robust is the Internet? Nature 406, 353–354.

87. Ma, L., Krishnan, R., and Montgomery, A.L. (2015). Latent Homophily or Social Influence? An Empirical Analysis of Purchase Within a Social Network. Manage. Sci. 61, 454–473.

88. Halberstam, Y., and Knight, B. (2016). Homophily, group size, and the diffusion of political information in social networks: Evidence from Twitter. J. Public Econ. 143, 73–88.

89. Mark, N.P. (2003). Culture and Competition: Homophily and Distancing Explanations for Cultural Niches. Am. Sociol. Rev. 68, 319–345.

90. Yavas, M., and Yuecel, G. (2013). Impact of homophily on diffusion dynamics over social networks. In ECMS 2013 Proceedings edited by: Webjorn Rekdalsbakken, Robin T. Bye, Houxiang Zhang (ECMS). 10.7148/2013-0888.

91. Borsook, D., Youssef, A.M., Simons, L., Elman, I., and Eccleston, C. (2018). When pain *gets stuck: the evolution of pain chronification and treatment resistance*. Pain 159, 2421–2436.

92. Gopalakrishnan, R., Burgess, R.C., Malone, D.A., Lempka, S.F., Gale, J.T., Floden, D.P., Baker, K.B., and Machado, A.G. (2018). Deep brain stimulation of the ventral striatal area for poststroke pain syndrome: a magnetoencephalography study. J. Neurophysiol. 119, 2118–2128.

93. García-Larrea, L., Peyron, R., Mertens, P., Gregoire, M.C., Lavenne, F., Le Bars, D., Convers, P., Mauguière, F., Sindou, M., and Laurent, B. (1999). Electrical stimulation of motor cortex for pain control: a combined PET-scan and electrophysiological study. Pain 83, 259–273.

94. Velasco, F., Carrillo-Ruiz, J.D., Castro, G., Argüelles, C., Velasco, A.L., Kassian, A., and Guevara, U. (2009). Motor cortex electrical stimulation applied to patients with complex regional pain syndrome. Pain 147, 91–98.

95. Duncan, G.H., Kupers, R.C., Marchand, S., Villemure, J.G., Gybels, J.M., and Bushnell, M.C. (1998). Stimulation of human thalamus for pain relief: possible modulatory circuits revealed by positron emission tomography. J. Neurophysiol. 80, 3326–3330.

96. Andy, O.J. (1983). Thalamic stimulation for chronic pain. Appl. Neurophysiol. 46, 116–123.

97. Rasche, D., Rinaldi, P.C., Young, R.F., and Tronnier, V.M. (2006). Deep brain stimulation for the treatment of various chronic pain syndromes. Neurosurg. Focus 21, E8.

98. Csárdi, G., and Nepusz, T. (2006). The igraph software package for complex network research. Complex Systems, 1695.

