## Supplementary material for "Graph theory analysis reveals an assortative pain network vulnerable to attacks": Table S1; Table S2; Table S3

**Table S1. Brain areas in the pain network.**

| Symbol | Brain area | Group | Group1 | Reference |
|---|---|---|---|---|
| ACAd | Anterior cingulate area, dorsal part | CH | Forebrain | [1–3] |
| ACAv | Anterior cingulate area, ventral part | CH | Forebrain | [4–6] |
| SSp | Primary somatosensory area | CH | Forebrain | [1,2] |
| SSs | Supplementary somatosensory area | CH | Forebrain | [1,2] |
| MOp | Primary motor area | CH | Forebrain | [1–3] |
| MOs | Secondary motor area | CH | Forebrain | [1–3] |
| AI | Agranular insular area | CH | Forebrain | [7–9] |
| RSPd | Retrosplenial area, dorsal part | CH | Forebrain | [10,11] |
| RSPv | Retrosplenial area, ventral part | CH | Forebrain | [10,11] |
| CA1 | Field CA1 | CH | Forebrain | [12,13] |
| CA3 | Field CA3 | CH | Forebrain | [12,13] |
| DG | Dentate gyrus | CH | Forebrain | [12,13] |
| CLA | Claustrum | CH | Forebrain | [12–14] |
| BLA | Basolateral amygdala nucleus | CH | Forebrain | [15–18] |
| CP | Caudoputamen | CH | Forebrain | [19,20] |
| ACB | Nucleus accumbens | CH | Forebrain | [21–23] |
| AAA | Anterior amygdala area | CH | Forebrain | [15,17,18] |
| CEA | Central amygdala nucleus | CH | Forebrain | [15,18,24,25] |
| VAL | Ventral anterior-lateral complex of the thalamus | BS | Forebrain | [26,27] |
| VM | Ventral medial nucleus of the thalamus | BS | Forebrain | [26,27] |
| VPL | Ventral posterolateral nucleus of the thalamus | BS | Forebrain | [26,27] |
| VPM | Ventral posteromedial nucleus of the thalamus | BS | Forebrain | [26,27] |
| LP | Lateral posterior nucleus of the thalamus | BS | Forebrain | [26,27] |
| PO | Posterior complex of the thalamus | BS | Forebrain | [26,27] |
| AV | Anteroventral nucleus of thalamus | BS | Forebrain | [26,27] |

| | | | | |
|---|---|---|---|---|
| AM | Anteromedial nucleus of thalamus | BS | Forebrain | [26,27] |
| LD | Lateral dorsal nucleus of thalamus | BS | Forebrain | [26,27] |
| MD | Mediodorsal nucleus of thalamus | BS | Forebrain | [26,27] |
| PVT | Paraventricular nucleus of the thalamus | BS | Forebrain | [28–30] |
| PT | Parataenial nucleus | BS | Forebrain | [26,27] |
| MH | Medial habenula | BS | Forebrain | [28] |
| LH | Lateral habenula | BS | Forebrain | [28,29] |
| ZI | Zona incerta | BS | Midbrain | [31–33] |
| PB | Parabrachial nucleus | BS | Midbrain | [34–36] |
| PG | Pontine gray | BS | Midbrain | |
| RN | Red nucleus | BS | Midbrain | [37,38] |
| RPO | Nucleus raphe pontis | BS | Midbrain | [37,38] |
| PAG | Periaqueductal gray | BS | Midbrain | [39] |
| CENT | Central lobule | CB | Hindbrain | [40,41] |
| CUL | Culmen | CB | Hindbrain | [40,41] |
| PYR | Pyramus (VIII) | CB | Hindbrain | [40,41] |
| NOD | Nodulus (X) | CB | Hindbrain | [40,41] |
| SIM | Simple lobule | CB | Hindbrain | [40,41] |
| AN | Ansiform lobule | CB | Hindbrain | [40,41] |
| PRM | Paramedian lobule | CB | Hindbrain | [40,41] |
| PFL | Paraflocculus | CB | Hindbrain | [40,41] |
| FN | Fastigial nucleus | CB | Hindbrain | [40,41] |
| IP | Interposed nucleus | CB | Hindbrain | [40,41] |

**Table S2. Centrality of all brain areas in the pain network.**

| Vertex | Degree | Closeness | Betweenness |
|---|---|---|---|
| ACAd | 45 | 0.17609827 | 1 |
| ACAv | 41 | 0.16582495 | 119 |
| SSp | 51 | 0.22318075 | 0 |
| SSs | 38 | 0.20510836 | 86 |
| MOp | 49 | 0.22807327 | 166 |
| MOs | 55 | 0.21569048 | 154 |
| AI | 39 | 0.16804397 | 93 |
| RSPd | 28 | 0.1711025 | 3 |
| RSPv | 37 | 0.15562876 | 40 |
| CA1 | 36 | 0.15403277 | 0 |
| CA3 | 21 | 0.18777668 | 82 |
| DG | 15 | 0.17941749 | 0 |
| CLA | 25 | 0.19679365 | 0 |
| BLA | 31 | 0.17118668 | 29 |
| CP | 58 | 0.21879034 | 216 |
| ACB | 34 | 0.19954181 | 129 |
| AAA | 17 | 0.19487944 | 24 |
| CEA | 36 | 0.22793885 | 318 |
| VAL | 30 | 0.17249759 | 12 |
| VM | 37 | 0.21124779 | 53 |
| VPL | 28 | 0.18454415 | 21 |
| VPM | 21 | 0.19529003 | 2 |
| LP | 37 | 0.18470254 | 32 |
| PO | 31 | 0.19807701 | 53 |
| AV | 22 | 0.12558275 | 0 |
| AM | 30 | 0.21195561 | 100 |

| | | | |
|---|---|---|---|
| LD | 25 | 0.16324199 | 45 |
| MD | 51 | 0.22575112 | 215 |
| PVT | 24 | 0.20913698 | 58 |
| PT | 18 | 0.15850886 | 57 |
| MH | 9 | 0.11750354 | 0 |
| LH | 15 | 0.18063207 | 0 |
| ZI | 50 | 0.20198961 | 315 |
| PB | 25 | 0.229578 | 730 |
| PG | 16 | 0.15370687 | 43 |
| RN | 24 | 0.171889 | 38 |
| RPO | 18 | 0.08327624 | 0 |
| PAG | 37 | 0.17434472 | 103 |
| CENT | 3 | 0.04748381 | 0 |
| CUL | 5 | 0.09123744 | 0 |
| PYR | 1 | 0.10607869 | 0 |
| NOD | 3 | 0.06280077 | 0 |
| SIM | 6 | 0.09244126 | 134 |
| AN | 7 | 0.09143168 | 112 |
| PRM | 4 | 0.11274039 | 0 |
| PFL | 8 | 0.09773672 | 1 |
| FN | 7 | 0.15933981 | 235 |
| IP | 8 | 0.08289321 | 294 |
| IO | 8 | 0.20393627 | 47 |

**Table S3. Community of each brain area in the pain network.**

| Vertex | Community No. |
|---|---|
| SIM | 1 |
| IP | 1 |
| AN | 2 |
| PRM | 2 |
| PFL | 2 |
| IO | 2 |
| ACAd | 3 |
| ACAv | 3 |
| MOs | 3 |
| AI | 3 |
| RSPd | 3 |
| RSPv | 3 |
| CLA | 3 |
| BLA | 3 |
| CP | 3 |
| AAA | 3 |
| CEA | 3 |
| VM | 3 |
| LP | 3 |
| AV | 3 |
| AM | 3 |
| LD | 3 |
| MD | 3 |
| ZI | 3 |

| | |
|---|---|
| RN | 3 |
| PAG | 3 |
| ACB | 4 |
| PVT | 4 |
| PT | 4 |
| RPO | 4 |
| CA1 | 5 |
| CA3 | 5 |
| DG | 5 |
| MH | 5 |
| LH | 5 |
| SSp | 6 |
| SSs | 6 |
| MOp | 6 |
| VAL | 6 |
| VPL | 6 |
| VPM | 6 |
| PO | 6 |
| PB | 7 |
| PG | 8 |
| CENT | 9 |
| CUL | 10 |
| PYR | 11 |
| NOD | 12 |
| FN | 13 |

Reference:


1. Vierck, C.J., Whitsel, B.L., Favorov, O.V., Brown, A.W., and Tommerdahl, M. (2013). Role of primary somatosensory cortex in the coding of pain. Pain *154*, 334–344.

2. Bushnell, M.C., Duncan, G.H., Hofbauer, R.K., Ha, B., Chen, J.I., and Carrier, B. (1999). Pain perception: is there a role for primary somatosensory cortex? Proc. Natl. Acad. Sci. U. S. A. *96*, 7705–7709.

3. Leite, J., Carvalho, S., Battistella, L.R., Caumo, W., and Fregni, F. (2017). Editorial: The Role of Primary Motor Cortex as a Marker and Modulator of Pain Control and Emotional-Affective Processing. Front. Hum. Neurosci. *11*, 270.

4. Hutchison, W.D., Davis, K.D., Lozano, A.M., Tasker, R.R., and Dostrovsky, J.O. (1999). Pain-related neurons in the human cingulate cortex. Nat. Neurosci. *2*, 403–405.

5. Fuchs, P.N., Peng, Y.B., Boyette-Davis, J.A., and Uhelski, M.L. (2014). The anterior cingulate cortex and pain processing. Front. Integr. Neurosci. *8*, 35.

6. Bliss, T.V.P., Collingridge, G.L., Kaang, B.-K., and Zhuo, M. (2016). Synaptic plasticity in the anterior cingulate cortex in acute and chronic pain. Nat. Rev. Neurosci. *17*, 485–496.

7. Starr, C.J., Sawaki, L., Wittenberg, G.F., Burdette, J.H., Oshiro, Y., Quevedo, A.S., and Coghill, R.C. (2009). Roles of the insular cortex in the modulation of pain: insights from brain lesions. J. Neurosci. *29*, 2684–2694.

8. Lu, C., Yang, T., Zhao, H., Zhang, M., Meng, F., Fu, H., Xie, Y., and Xu, H. (2016). Insular Cortex is Critical for the Perception, Modulation, and Chronification of Pain. Neurosci. Bull. *32*, 191–201.

9. Mutschler, I., Ball, T., Wankerl, J., and Strigo, I.A. (2012). Pain and emotion in the insular cortex: evidence for functional reorganization in major depression. Neurosci. Lett. *520*, 204–209.

10. Barrière, D.A., Hamieh, A.M., Magalhães, R., Traoré, A., Barbier, J., Bonny, J.-M., Ardid, D., Busserolles, J., Mériaux, S., and Marchand, F. (2019). Structural and functional alterations in the retrosplenial cortex following neuropathic pain. Pain *160*, 2241–2254.

11. Wik, G., Fischer, H., Finer, B., Bragee, B., Kristianson, M., and Fredrikson, M. (2006). Retrospenial cortical deactivation during painful stimulation of fibromyalgic patients. Int. J. Neurosci. *116*, 1–8.

12. Mutso, A.A., Radzicki, D., Baliki, M.N., Huang, L., Banisadr, G., Centeno, M.V., Radulovic, J., Martina, M., Miller, R.J., and Apkarian, A.V. (2012). Abnormalities in hippocampal functioning with persistent pain. J. Neurosci. *32*, 5747–5756.

13. Grilli, M. (2017). Chronic pain and adult hippocampal neurogenesis: translational implications from preclinical studies. J. Pain Res. *10*, 2281–2286.

14. Gracely, R.H., Geisser, M.E., Giesecke, T., Grant, M.A.B., Petzke, F., Williams, D.A., and Clauw, D.J. (2004). Pain catastrophizing and neural responses to pain among persons with fibromyalgia. Brain *127*, 835–843.

15. Neugebauer, V. (2015). Amygdala pain mechanisms. Handb. Exp. Pharmacol. *227*, 261–284.

16. Corder, G., Ahanonu, B., Grewe, B.F., Wang, D., Schnitzer, M.J., and Scherrer, G. (2019). An amygdalar neural ensemble that encodes the unpleasantness of pain. Science *363*, 276–281.

17. Li, Z., Wang, J., Chen, L., Zhang, M., and Wan, Y. (2013). Basolateral amygdala lesion inhibits the development of pain chronicity in neuropathic pain rats. PLoS One *8*, e70921.

18. Thompson, J.M., and Neugebauer, V. (2017). Amygdala Plasticity and Pain. Pain Res. Manag. *2017*, 8296501.

19. Barceló, A.C., Filippini, B., and Pazo, J.H. (2012). The striatum and pain modulation. Cell. Mol. Neurobiol. *32*, 1–12.

20. Borsook, D., Upadhyay, J., Chudler, E.H., and Becerra, L. (2010). A key role of the basal ganglia in pain and analgesia--insights gained through human functional imaging. Mol. Pain *6*, 27.

21. Baliki, M.N., Geha, P.Y., Fields, H.L., and Apkarian, A.V. (2010). Predicting value of pain and analgesia: nucleus accumbens response to noxious stimuli changes in the presence of chronic pain. Neuron *66*, 149–160.

22. Harris, H.N., and Peng, Y.B. (2020). Evidence and explanation for the involvement of the nucleus accumbens in pain processing. Neural Regeneration Res. *15*, 597–605.

23. DosSantos, M.F., Moura, B. de S., and DaSilva, A.F. (2017). Reward Circuitry Plasticity in Pain Perception and Modulation. Front. Pharmacol. *8*, 790.

24. Warlow, S.M., Naffziger, E.E., and Berridge, K.C. (2020). The central amygdala recruits mesocorticolimbic circuitry for pursuit of reward or pain. Nat. Commun. *11*, 2716.

25. O'Neill, P.-K., and Meszaros, J. (2020). Chronic Pain Releases Parabrachial Activity from Central Amygdala Inhibition. J. Neurosci. *40*, 7996–7998.

26. Gustin, S.M., Peck, C.C., Wilcox, S.L., Nash, P.G., Murray, G.M., and Henderson, L.A. (2011). Different pain, different brain: thalamic anatomy in neuropathic and non-neuropathic chronic pain syndromes. J. Neurosci. *31*, 5956–5964.

27. Sotgiu, M.L. (2001). The Thalamus and Pain. In Neuroscience: Focus on Acute and Chronic Pain (Springer Milan), pp. 37–42.

28. Shelton, L., Becerra, L., and Borsook, D. (2012). Unmasking the mysteries of the habenula in pain and analgesia. Prog. Neurobiol. *96*, 208–219.

29. Shelton, L., Pendse, G., Maleki, N., Moulton, E.A., Lebel, A., Becerra, L., and Borsook, D. (2012). Mapping pain activation and connectivity of the human habenula. J. Neurophysiol. *107*, 2633–2648.

30. Kang, S., Li, J., Zuo, W., Chen, P., Gregor, D., Fu, R., Han, X., Bekker, A., and Ye, J.-H. (2019). Downregulation of M-channels in lateral habenula mediates hyperalgesia during alcohol withdrawal in rats. Sci. Rep. *9*, 2714.

31. Lu, C.W., Harper, D.E., Askari, A., Willsey, M.S., Vu, P.P., Schrepf, A.D., Harte, S.E., and Patil, P.G. (2021). Stimulation of zona incerta selectively modulates pain in humans. Sci. Rep. *11*, 8924.

32. Moon, H.C., and Park, Y.S. (2017). Reduced GABAergic neuronal activity in zona incerta causes neuropathic pain in a rat sciatic nerve chronic constriction injury model. J. Pain Res. *10*, 1125–1134.

33. Masri, R., Quiton, R.L., Lucas, J.M., Murray, P.D., Thompson, S.M., and Keller, A. (2009). Zona incerta: a role in central pain. J. Neurophysiol. *102*, 181–191.

34. Chiang, M.C., Bowen, A., Schier, L.A., Tupone, D., Uddin, O., and Heinricher, M.M. (2019). Parabrachial Complex: A Hub for Pain and Aversion. J. Neurosci. *39*, 8225–8230.

35. Roeder, Z., Chen, Q., Davis, S., Carlson, J.D., Tupone, D., and Heinricher, M.M. (2016). Parabrachial complex links pain transmission to descending pain modulation. Pain *157*, 2697–2708.

36. Palmiter, R.D. (2018). The Parabrachial Nucleus: CGRP Neurons Function as a General Alarm. Trends Neurosci. *41*, 280–293.

37. Yang, Q.-Q., Li, H.-N., Zhang, S.-T., Yu, Y.-L., Wei, W., Zhang, X., Wang, J.-Y., and Zeng, X.-Y. (2020). Red nucleus IL-6 mediates the maintenance of neuropathic pain by inducing the productions of TNF-α and IL-1β through the JAK2/STAT3 and ERK signaling pathways. Neuropathology. 10.1111/neup.12653.

38. François, A., Low, S.A., Sypek, E.I., Christensen, A.J., Sotoudeh, C., Beier, K.T., Ramakrishnan, C., Ritola, K.D., Sharif-Naeini, R., Deisseroth, K., et al. (2017). A Brainstem-Spinal Cord Inhibitory Circuit for Mechanical Pain Modulation by GABA and Enkephalins. Neuron *93*, 822–839.e6.

39. Mokhtar, M., and Singh, P. (2020). Neuroanatomy, Periaqueductal Gray. In StatPearls (StatPearls Publishing).

40. Moulton, E.A., Schmahmann, J.D., Becerra, L., and Borsook, D. (2010). The cerebellum and pain: passive integrator or active participator? Brain Res. Rev. *65*, 14–27.

41. Coombes, S.A., and Misra, G. (2016). Pain and motor processing in the human cerebellum. Pain *157*, 117–127.
